# Orthologous transcription factor replacement reveals that stable TFIIIC complexes are required for proper mitotic chromosome segregation

**DOI:** 10.1101/2025.01.30.635671

**Authors:** Akshi Gupta, Po-Chen Hsu, Richard Ron R. Litan, Jun-Yi Leu

**Affiliations:** Molecular and Cell Biology, Taiwan International Graduate Program, Institute of Molecular Biology, Academia Sinica and Graduate Institute of Life Sciences, National Defense Medical Center, Taipei, Taiwan; Institute of Molecular Biology, Academia Sinica, Taipei, 11529, Taiwan

**Keywords:** essential gene, genetic incompatibility, transcription factor, TFIIIC, cohesin, tRNA, evolution

## Abstract

Transcription factors are speculated to play crucial roles in adaptive evolution. Using ortholog replacement of essential transcription factors (eTFs) from other yeast species, we investigated how eTFs can change. Several orthologs could not fully complement *Saccharomyces cerevisiae* mutants, indicating that functions or interactions of these eTFs have changed, rendering them incompatible. We further characterized TFIIIC, a fast-evolving protein complex that assists RNA polymerase III-mediated transcription, which exhibited complete or partial incompatibility in several subunits. In the orthologous Tfc7-replacement line, binding of TFIIIC to tRNA genes was reduced, yet tRNA abundance was not severely affected. However, the chromosomes of Tfc7-replacement cells were often mis-segregated during mitosis and their fitness was further reduced in a spindle checkpoint mutant. Our chromatin-immunoprecipitation experiments uncovered that unstable TFIIIC binding results in defective cohesion loading, leading to chromosome mis-segregation. Swapping the highly divergent C-terminal domain of Tfc7 orthologs rescued its interaction with Tfc1 and cell fitness, supporting that incompatibility is caused by altered interactions between complex subunits. Our results reveal distinct essential functions of a well-studied protein complex.

## Introduction

Essential genes are those required for cell survival under optimal growth conditions and they are often involved in basic but fundamental cellular processes. In general, the protein sequences of essential genes are more conserved than those of non-essential genes (Hirsh and Fraser, 2001), and essential genes are more likely to be retained in the genome during species divergence (Heinicke et al., 2007; Kim et al., 2010). Nonetheless, recent studies have shown that essential genes can have very different evolutionary trajectories (Lai et al., 2023). Moreover, some yeast essential genes can be replaced by bacterial orthologs, whereas others cannot even be exchanged between different yeast species (Kachroo et al., 2017; Kachroo et al., 2015; Lai et al., 2023). Thus, apart from their well-known essential functions, the evolution of essential genes can be influenced by other unidentified factors, such as pleiotropic effects or moonlighting functions.

Essential genes involved in transcriptional regulation represent an interesting functional group. Transcription regulatory networks allow cells to alter their physiology in response to external or internal signals. Elements regulating gene expression include DNA methylation, histone modification, chromatin remodeling, and transcriptional activators/repressors (Isbel et al., 2022). Proteins involved in these processes are broadly classified as transcription regulatory proteins or transcription factors (TFs). Altered transcriptional regulation has been observed in many adaptive evolutionary events, such as enhanced heavy metal tolerance in microorganisms, changed body plans in animals, and variation of flowering patterns in plants (Chang et al., 2013; Chang and Leu, 2011; Della Pina et al., 2014; Peter and Davidson, 2011; Wray, 2007). Moreover, increased numbers and functional diversification of TFs are speculated to be crucial factors contributing to the evolution of organismal complexity (Cheatle Jarvela and Hinman, 2015; Molina and van Nimwegen, 2008). Recent studies have shown that TFs can perform functions other than transcriptional regulation, such as nuclear or chromosomal organization (Andersson and Tracey, 2011; Morgan and Shilatifard, 2023; Yang et al., 2022). These pleiotropic functions may add to the complexity of TF evolution.

In the budding yeast *Saccharomyces cerevisiae*, 6.6% (437/6611) of the genome encodes TFs, with 93 of those TFs being essential (Cherry et al., 2012; Rossi et al., 2021; Sorrells et al., 2018; Teixeira et al., 2022). We adopted a gene ortholog replacement strategy to investigate some of these essential TFs (eTFs). Ortholog replacement represents a useful approach for investigating the evolution of essential genes (Kachroo et al., 2017; Kachroo et al., 2015; Lai et al., 2023). If a replaced ortholog sourced from a different species cannot fully rescue the mutant fitness of its *S. cerevisiae* host, then the function or interactions of that orthologous protein have likely changed in its parental species. The incompatible TF orthologs may be further dissected to understand how the changes occurred.

In our previous study (Lai et al., 2023), we tested orthologs from four ascomycete yeast species, namely *Naumovozyma castellii*, *Kluyveromyces lactis*, *Yarrowia lipolytica*, and *Schizosaccharomyces pombe*, which are estimated to have diverged from their common ancestor with *S. cerevisiae* about 50, 100, 270 and 420 million years ago, respectively (Galagan et al., 2005; Taylor and Berbee, 2006). Most of the essential gene orthologs from *N. castellii* or *K. lactis* we tested were compatible with *S. cerevisiae*, except for a few fast-evolving ones (Lai et al., 2023). In the current study, we were interested in exploring this fast-evolving pattern, so we performed gene replacement using orthologs from *N. castellii* and *K. lactis* to assess if we could identify fast-evolving eTFs (Fig. 1A). Moreover, since *K. lactis* diverged from *S. cerevisiae* before an ancient whole genome duplication (WGD) event and *N. castellii* diverged after the WGD (Taylor and Berbee, 2006), gene evolution in these two species might be driven by different forces influenced by genome duplication.

**Figure 1.**
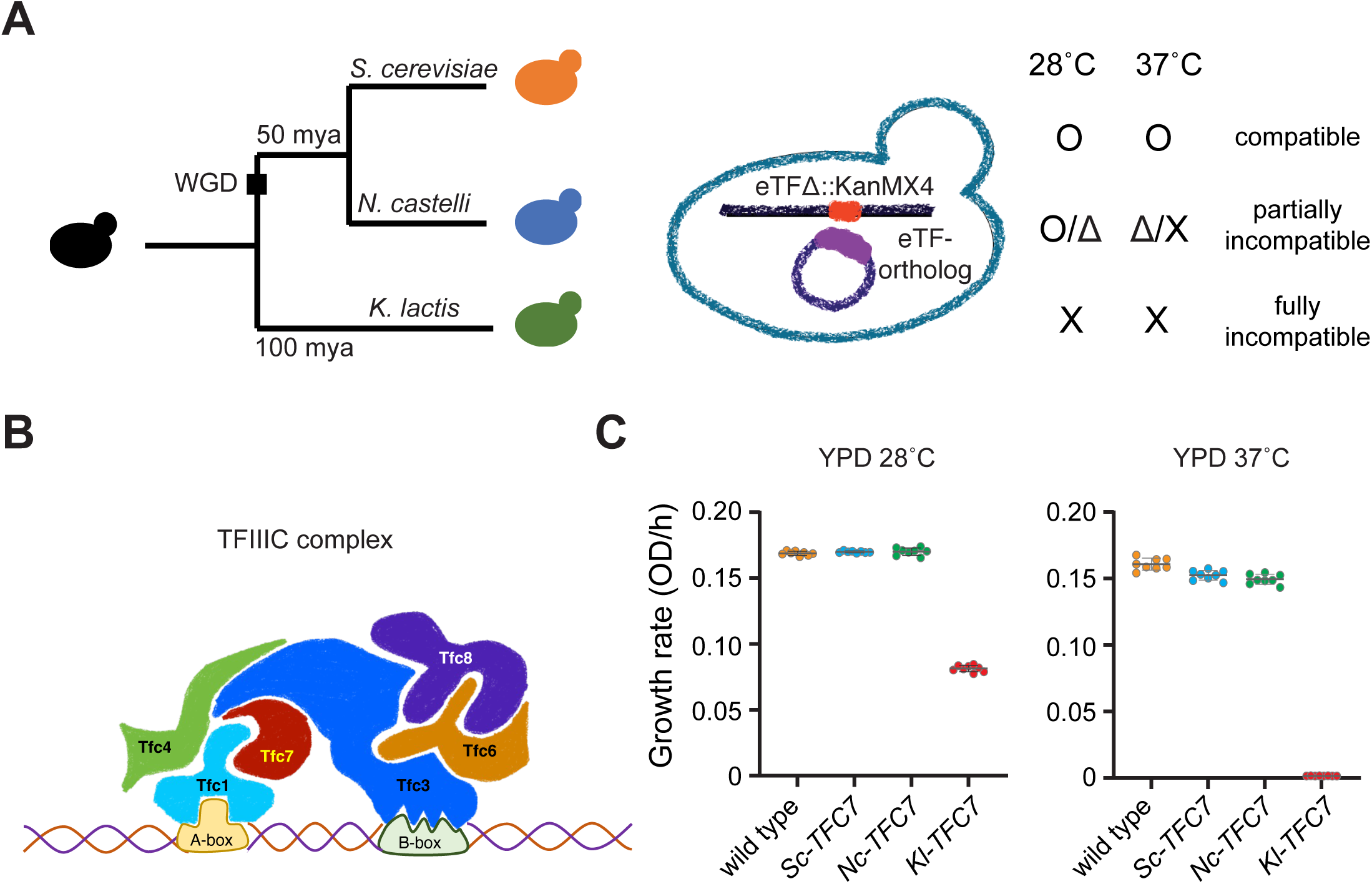
Ortholog replacement assays reveal incompatibility between essential TFs and the *S. cerevisiae* background. (A) Simplified phylogenetic relationship of *S. cerevisiae, N. castellii* and *K. lactis* showing the estimated time of divergence. In the haploid replacement line, the native eTF gene was disrupted by inserting a drug cassette, and the *N. castellii* or *K. lactis* orthologous gene carried by a plasmid was tested to see how much it could complement the essential function. Yeast cells were grown on rich medium plates (YPD) at 28 °C (optimal temperature) or 37 °C (heat stress) to examine ortholog compatibility. O, normal growth; Δ, slow growth; X, no growth. (B) Schematic diagram of the TFIIIC complex, as interpreted from multiple sources (Male et al., 2015; Vorländer et al., 2020), showing the Tfc7 subunit interacting with other subunits of the TFIIIC complex. The complex binds to the intrinsic promoter sequences, known as the A-box and B-box. (C) The replacement line carrying *Kl-TFC7* exhibits growth defects at 28 °C and no growth at 37 °C. Yeast cells carrying different plasmids were grown in YPD liquid medium, and their growth rates were measured by the plate reader. Eight replicates were measured in this experiment. The mean and standard error are shown.

While testing the essential TFs, we observed complete incompatibility in the replacement line when orthologous Tfc8 was derived from *K. lactis*. Tfc8 is a subunit of the RNA polymerase III transcription initiation factor C (TFIIIC) complex; a complex conserved from yeast to humans and with the well-established primary function of facilitating transcription of RNA polymerase III (Deprez et al., 1999; Male et al., 2015). RNA polymerase III is responsible for the synthesis of tRNAs, 5S rRNA, and other non-coding RNAs (ncRNAs) in eukaryotic cells (Dieci et al., 2007; White, 2011). TFIIIC comprises two subcomplexes, τA and τB, that bind two gene internal DNA motifs, i.e., A-box and B-box (Galli et al., 1981; Marzouki et al., 1986). τA is composed of τ131 (Tfc4), τ95 (Tfc1), and τ55 (Tfc7), whereas τB comprises τ138 (Tfc3), τ91 (Tfc6), and τ60 (Tfc8) (Male et al., 2015; Soragni and Kassavetis, 2008). All six subunits of the TFIIIC complex are essential for cell viability in yeast and are responsible for recruiting TFIIIB and RNA polymerase III to tRNA for transcription (Male et al., 2015).

Previous studies have shown that, in addition to the transcription machinery, chromosomal regions encoding tRNAs are bound by other chromatin proteins, including the architectural SMC proteins, nuclear pore proteins, chromatin remodelers, and histone modifiers (D’Ambrosio et al., 2008a; Dhillon et al., 2009; Donze, 2012; Glynn et al., 2004; Oki and Kamakaka, 2005). Nonetheless, the relationships between these proteins remain to be fully elucidated in most cases. By analyzing the cellular defects caused by incompatible TFIIIC subunits, we have discovered that an unstable hybrid TFIIIC complex does not severely affect tRNA abundance, yet it has a strong impact on cohesin loading and thus proper mitotic chromatid segregation. Together, our results reveal that eTFs can have essential secondary functions, which may be overlooked due to their more obvious primary functions. Moreover, the multi-functionality of a protein may impact its evolutionary trajectory.

## Results

### Essential transcription factors exhibit partial or complete incompatibility in ortholog replacement lines

To identify fast-evolving eTFs, we constructed *S. cerevisiae* ortholog replacement lines using orthologs from *N. castellii* and *K. lactis* (Fig. 1A). Promoter function can change rapidly, even among closely related species (Borneman et al., 2007; Lai et al., 2023; Tirosh et al., 2009), but alterations in essential protein functions or interactions have been less well explored. In this study, we focused solely on protein evolution. We selected four eTF genes involved in various categories of transcriptional regulation for our replacement experiments, namely *FHL1*, *PDC2*, *RSC4*, and *TFC8* (Supplemental Table S1 and Fig. S1). The respective encoded TFs operate in transcriptional regulation of ribosomal protein genes (Fhl1) and thiamine synthesis genes (Pdc2), chromatin remodeling (Rsc4), and RNA polymerase III transcriptional initiation (Tfc8), respectively. Expression of the *S. cerevisiae* genes and their orthologs was driven by the strong Tet-Off promoter (see Materials and Methods) to ensure constitutive gene expression. Successful replacement lines were grown at 28 °C and 37 °C to assay compatibility. The latter 37 °C stress condition allowed us to detect partial incompatibility since a growth defect caused by partial incompatibility is often more apparent at high temperatures.

For *PDC2*, all replacement lines behaved similarly to the cells carrying *S. cerevisiae-*derived *PDC2* (*Sc-PDC2*), supporting that it is an evolutionarily static TF (Supplemental Table S1 and Fig. S2A). In contrast, the other gene replacement lines showed partial or complete incompatibility from at least one ortholog (Supplemental Table S1 and Fig. S2A). These data indicate that the interactions or functions of some eTFs have been altered between *S. cerevisiae* and the other two yeast species. Among the tested TFs, the *K. lactis-TFC8* (*Kl-TFC8*) line was the only one exhibiting complete incompatibility, since cells carrying *Kl-TFC8* were inviable.

### Several essential Transcription Factor III C (TFIIIC) subunits exhibit similar evolutionary patterns

Tfc8 is a subunit of the TFIIIC complex that works together with five other essential subunits to recruit RNA polymerase III transcription factor B (TFIIIB) and to help RNA polymerase III to transcribe its target genes (Deprez et al., 1999). Our previous study showed that some protein complex subunits co-evolve and exhibit similar evolving patterns (Lai et al., 2023). Since Kl-Tfc8 proved completely incompatible in the *S. cerevisiae* background, we examined the ortholog compatibility of other TFIIIC subunits, i.e., Tfc1, Tfc4, Tfc6 and Tfc7 (Fig. 1B) (Arrebola et al., 1998; Male et al., 2015). Tfc3 was not included since we could not obtain a deletion strain despite several trials. The *TFC3* gene is located close to the centromere of Chromosome I, which may explain our inability to delete the gene by homologous recombination.

Interestingly, only *N. castellii*-*TFC1* (*Nc*-*TFC1*) and *Nc-TFC7* cells grew similarly to wild-type cells. *Nc-TFC4* and *Nc-TFC6* cells grew slowly and were sensitive to high temperatures. *Kl-TFC1*, *Kl-TFC4*, and *Kl-TFC6* were all completely incompatible with *S. cerevisiae*, i.e., similar to the outcome for *Kl-TFC8.* The *Kl-TFC7* replacement line grew slowly at 28 °C but became inviable at 37 °C (Fig. 1C and Supplemental Fig. S2B), revealing partial incompatibility. These results indicate that even within a protein complex, subunits can exhibit different levels of compatibility, indicating the formation and possible co-evolution of subcomplexes within a larger protein complex. In order to understand the molecular basis of TFIIIC incompatibility, we chose the slow-growing *Kl-TFC7* replacement line for further investigation.

### tRNA levels are not severely affected in *Kl-TFC7* cells

Since the primary known function of TFIIIC is to facilitate RNA polymerase III transcription of tRNAs (Deprez et al., 1999; Male et al., 2015), first we examined levels of total tRNAs. There are 286 tRNA genes spread across the nuclear genome of *S. cerevisiae*, with multiple copies for each amino acid codon required for translation of mRNAs into proteins (Chan et al., 2021). Interestingly, we detected similar levels of total tRNAs and 5S rRNA in both the replacement line and wild-type cells (Fig. 2A), indicating that the incompatibility of *Kl-TFC7* does not severely impact steady-state levels of total tRNAs.

**Figure 2.**
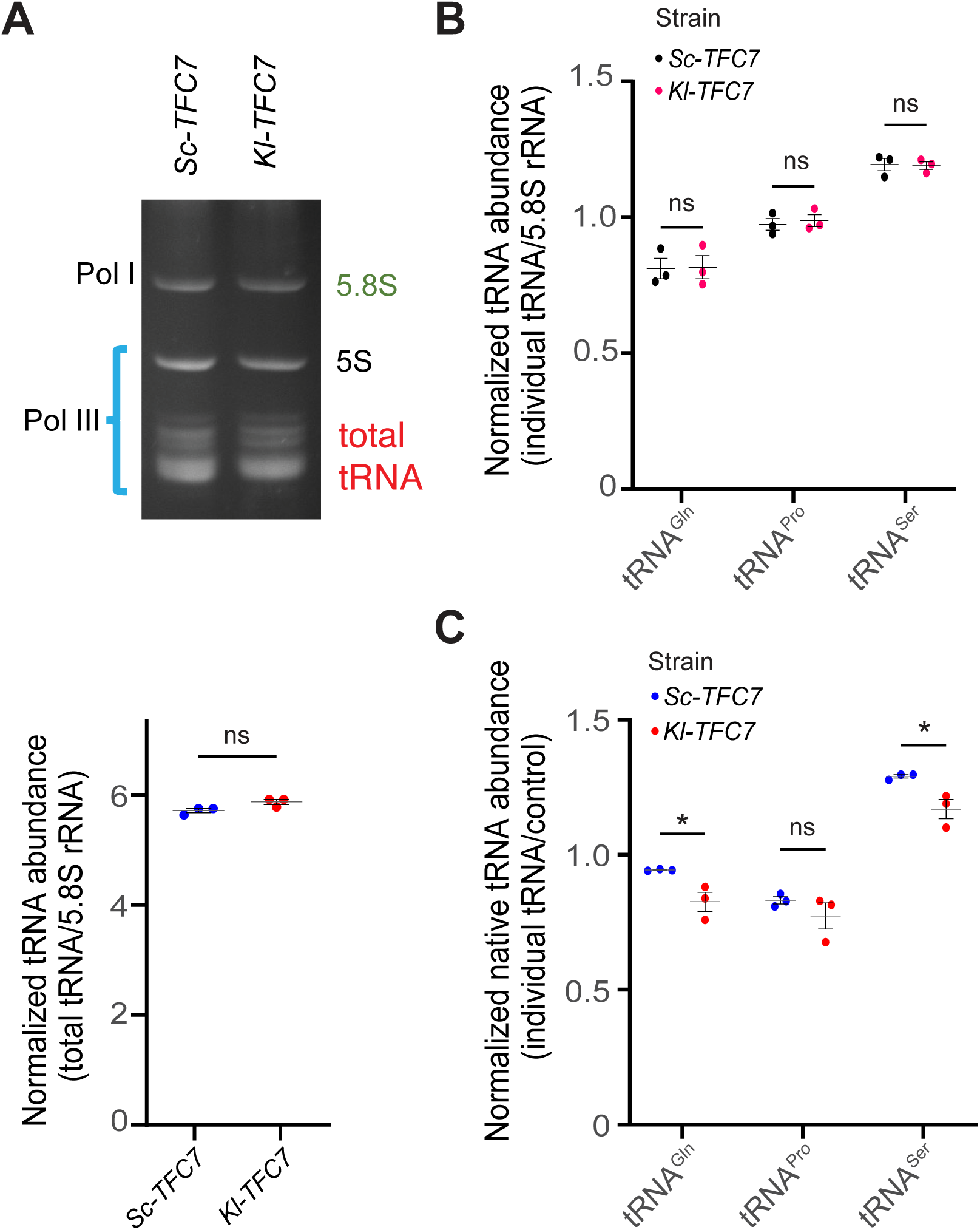
tRNA transcription is not severely compromised in *Kl-TFC7* cells. (A) *Kl-TFC7* cells have a total tRNA level similar to *Sc-TFC7* cells. Total RNA was run on the 12% TBE urea gel and the signal intensity of total tRNA was normalized to the intensity of 5.8S rRNA, which is transcribed by RNA polymerase I. The bottom panel shows the data of three replicates. ns, p-value > 0.05 (Student’s t-test, n = 3). (B) q-PCR of small RNA isolated from *Kl-TFC7* and *Sc-TFC7* cells, revealing no significant difference in the abundances of tRNAs for glutamine, proline and serine. ns, p-value > 0.05 (Student’s t-test, n = 3). (C) q-PCR of nascent RNA isolated from *Kl-TFC7* and *Sc-TFC7* cells reveals slight differences in the abundances of tRNAs for glutamine and serine, yet no difference in the tRNA for proline. In this experiment, *S. pombe* native tubulin RNA was used as a spike-in internal control. The mean and standard error are shown. ns, p-value > 0.05; *, p < 0.05 (Student’s t-test, n = 3).

In addition to total tRNA abundance, we also measured a few individual tRNAs by quantitative PCR (q-PCR), including tRNA^Gln^(TTG/CTG), tRNA^Pro^(AGG/GGG/TGG/CGG), and tRNA^Ser^(CGA). In the budding yeast genome, there are 10, 11, and 17 copies of tRNA^Gln^, tRNA^Pro^, and tRNA^Ser^, respectively, with our primer sets recognizing all tRNA^Gln^, 9 copies of tRNA^Pro^, and 11 copies of tRNA^Ser^. Our q-PCR results revealed no differences in the levels of these tRNAs between the *Sc-TFC7* and *Kl-TFC7* lines (Fig. 2B).

tRNAs display highly stable secondary structures, along with longer half-lives compared to most mRNAs found in cells (Chan et al., 2018; Lorenz et al., 2017). The average half-life of mRNAs in budding yeast cells is close to 4.8 minutes *in vivo* (Chan et al., 2018), whereas tRNAs have an average half-life close to 100 hours and with a high turnover capacity (Lorenz et al., 2017). To establish if newly transcribed tRNAs are affected in *Kl-TFC7* cells, we purified nascent tRNAs (see Materials and Methods) and quantified the same set of individual tRNAs described above by means of q-PCR. The *Kl-TFC7* cells only presented a slightly lower level of native tRNAs than the *Sc-TFC7* control cells (Fig. 2C). Thus, our data indicate that the growth defect of *Kl-TFC7* cells is probably not attributable to reduced tRNA levels.

### Sister chromatids are mis-segregated during mitosis in *Kl-TFC7* cells

The *Kl-TFC7* replacement line grew more slowly than the *Sc-TFC7* control line under normal conditions (Fig. 1C). During cell growth, cell cycle checkpoints monitor many cellular processes. When problems are detected, they slow down or arrest the cell cycle to repair the defects. For example, during mitosis, duplicated sister chromatids must be held together by cohesin and attached to the bipolar spindle to generate spindle tension. If sister chromatids fail to generate such tension, it triggers the spindle checkpoint to block precocious onset of anaphase (Musacchio and Hardwick, 2002). To investigate if the slow growth of *Kl-TFC7* cells was caused by checkpoint events, we deleted *MAD2,* encoding a major regulator of the spindle checkpoint. In the absence of Mad2 protein, cells proceed through mitosis irrespective of whether all sister kinetochores are under spindle tension (Li and Murray, 1991). We found that *MAD2* deletion had only a negligible effect on *Sc-TFC7* cells because sister chromatids in wild-type cells were held together by cohesin to generate proper tension during mitosis. However, the fitness of *Kl-TFC7* cells was severely reduced when *MAD2* was deleted (Fig. 3A and 3B). This outcome indicates that the incompatible Kl-Tfc7 protein may cause loss of sister chromatid cohesion or misorientation of sister kinetochores, thereby promoting frequent chromosome mis-segregation and ultimately cell death if Mad2 is absent.

**Figure 3.**
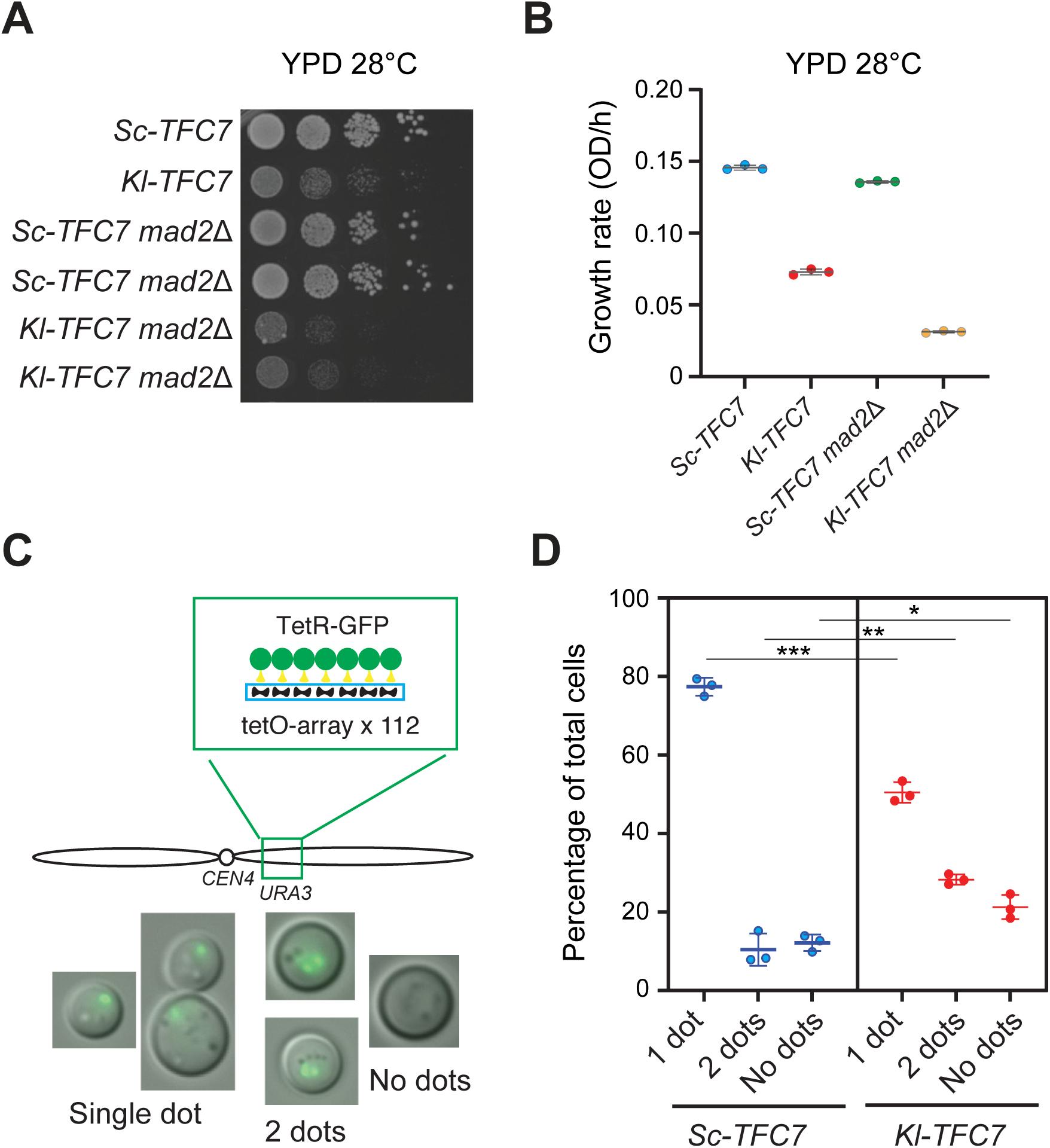
*Kl-TFC7* cells exhibit chromosome segregation defects. (A) Deleting the spindle checkpoint gene *MAD2* enhances the growth defect of *Kl-TFC7* cells on solid medium. Spot assay of various lines at 28 °C revealed significantly impaired growth for *Kl-TFC7 mad2*Δ cells compared to *Kl-TFC7* cells. (B) *MAD2* deletion enhances the growth defect of *Kl-TFC7* cells in liquid medium. Cells were grown in YPD liquid medium at 28 °C and growth rates were measured using plate readers. Three replicates were measured in this experiment. (C) A TetR-GFP system for monitoring chromosome segregation. To quantify the chromosome segregation defect, a TetO array of 112 repeats was first inserted into the right arm of Chromosome IV. When cells expressed TetR-GFP with a nuclear localization signal, TetR-GFP recognized the TetO array and presented a strong GFP signal under microscopy. Since our strains are haploid, normal cells should only present one GFP dot. Cells with two GFP dots indicate sister chromatid cohesion defects or chromosome mis-segregation. Cells with no GFP dots represent aneuploid or dead cells. (D) *Kl-TFC7* cells show a significant increase in cells with two GFP dots or no dots. In each replicate, at least 400 cells were counted. The mean and standard error are shown. *, p-value < 0.05; **, p < 0.01; ***, p < 0.005 (Student’s t-test, n = 3).

To further examine the chromosome segregation defect of *Kl-TFC7* cells, we engineered insertion of a tetO repeat array into chromosome IV that would be bound by TetR-GFP and allow us to monitor by GFP-dot assay chromosome segregation under microscopy (Chen, 2019; Fuchs et al., 2002). Normal haploid cells should be observed as only containing one GFP dot (representing sister chromatids) even after DNA replication since the sister chromatids are held together by cohesin. Cells with two GFP dots are either aneuploid cells resulting from chromosome mis-segregation or G2-phase cells with defects in sister chromatid cohesion. Cells with no GFP dot represent aneuploid cells lacking the labeled chromosome or they are dead cells (Fig. 3C). Only a small proportion of cells from the *Sc-TFC7* line presented two or no GFP dots. In contrast, 29% of cells of the *Kl-TFC7* line presented two GFP dots and 20% had no dots (Fig. 3D). This data is consistent with our findings on *mad2* deletion, indicating that chromosome mis-segregation is the primary defect of the *Kl-TFC7* line.

### Replacing Kl-Tfc7 reduces the association of the TFIIIC complex with chromatin

The TFIIIC complex binds to DNA to facilitate RNA pol III transcription and several aspects of chromosomal organization such as tRNA nucleolar clustering, the peripheral nuclear positioning of extra TFIIIC (ETC) sites, condensin loading, and mating locus silencing (D’Ambrosio et al., 2008b; Deprez et al., 1999; Haeusler et al., 2008; Hamdani et al., 2019a; Hiraga et al., 2012). To determine if the DNA binding of TFIIIC is affected by replacement with Kl-Tfc7, we performed chromatin immunoprecipitation (ChIP) sequencing by pulling down a tagged endogenous Tfc3 protein. Three biological replicates were sequenced for each cell type and only sites with binding peaks in all three replicates were considered as enriched sites to control the accuracy of target sequence identification (Supplemental Table S2).

In total, we identified 446 DNA binding sites in both *Sc-TFC7* and *Kl-TFC7* cells. Consistent with previous studies (Eaton et al., 2010; Hiraga et al., 2012; Simms et al., 2008), the TFIIIC complex stably bound to most tRNA genes, ETC sites, and other non-coding transcription sites in the *Sc-TFC7* cells (Fig. 4A and Supplemental Table S2). In contrast, many of the DNA binding peaks in *Kl-TFC7* cells were weaker than those of *Sc-TFC7* cells, especially for the tRNA genes (Fig. 4B and Supplemental Fig. S3).

**Figure 4.**
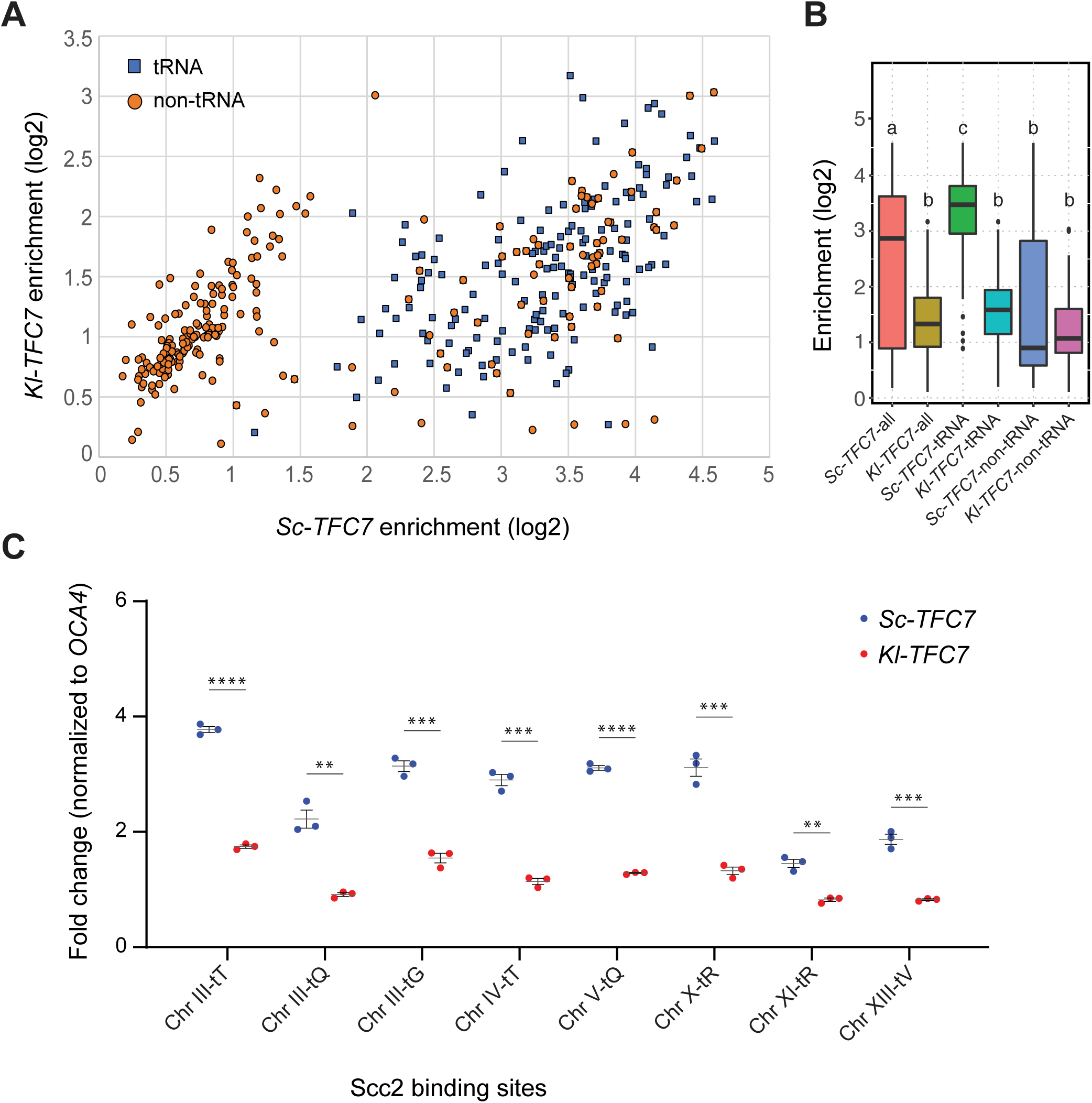
*Kl-TFC7* cells display low target binding of TFIIIC complexes and cohesin loaders. (A) Scatter plot from Tfc3 chromatin immunoprecipitation (ChIP) sequencing data. The enrichment scores (log10) of individual Tfc3 binding sites in *Sc-TFC7* and *Kl-TFC7* cells are indicated on the x-axis and y-axis, respectively. (B) Binding of Tfc3 to tRNA sites is significantly reduced in *Kl-TFC7* cells. Boxplots show the enrichment scores of Tfc3 binding to all targets, tRNA sites, and non-tRNA sites. Data are derived from Tfc3 ChIP-seq experiments. Different letters above each boxplot indicate statistically significant differences (ANOVA and Tukey’s HSD post-hoc test, adjusted p-value < 0.05). (C) Scc2 DNA binding is significantly reduced in *Kl-TFC7* cells. ChIP experiments were conducted by pulling down Scc2-13Myc, and eight known Scc2 binding sites were examined using q-PCR. The *OCA4* locus was used as an internal control since Scc2 does not bind to this site (Hamdani et al., 2019b). The mean and standard error are shown. **, p < 0.01; ***, p < 0.005; ****, p < 0.001 (Student’s t-test, n = 3).

Although our ChIP data revealed that binding of TFIIIC to tRNA genes was strongly reduced in the *Kl-TFC7* line, we have already shown that the abundance of tRNAs (both in total and for several individual ones) was not significantly affected (Fig. 2). However, we also observed a more severe fitness defect caused by chromosome mis-segregation in *Kl-TFC7* cells (Fig. 3). Previous studies have shown that condensin-binding sites overlap with TFIIIC binding sites, and condensin subunits often colocalize with the cohesin loaders Scc2 and Scc4 (D’Ambrosio et al., 2008b; Haeusler et al., 2008), raising the possibility that cohesin loading is compromised due to unstable binding of TFIIIC in the *Kl-TFC7* line, thereby leading to chromosome mis-segregation. We constructed strains carrying TAP-tagged or Myc-tagged cohesin subunits and loading factors, i.e., Smc1, Smc3, Mcd1 (Scc1), Irr1 (Scc3), and Scc2. All of the tagged strains showed significantly reduced fitness in the *Kl-TFC7* background (Supplemental Fig. S4), revealing a synthetic sickness effect and indicating a possible interaction between cohesin and TFIIIC. To examine if cohesin loading was affected in the *Kl-TFC7* line, we used the least sick Scc2 Myc-tagged strain to perform ChIP experiments and to quantify by q-PCR binding of Scc2 to a few tRNA loci. We observed significantly reduced binding for all loci (Fig. 4C), providing direct evidence that *Kl-TFC7* cells display cohesin loading defects.

### Kl-Tfc7 fails to interact properly with Sc-Tfc1

This reduced ChIP peak intensity (Fig. 4C) might result from reduced total TFIIIC abundance or/and unstable binding. Western blots showed that Tfc3 protein abundance in *Kl-TFC7* cells was reduced to about 25% the level of *Sc-TFC7* cells (Fig. 5A). Since the tRNA binding signal intensities of Tfc3 in *Kl-TFC7* are decreased ∼4-fold relative to those in *Sc-TFC7* (Fig. 4B), the reduced abundance of TFIIIC is possibly the primary reason for this outcome. However, the *TFC3* gene in *Kl-TFC7* cells is exactly the same as that in *Sc-TFC7* cells. Consequently, why is Tfc3 abundance affected by the introduction of *Kl-TFC7*? Some protein complex subunits are degraded by the proteostatic machinery if they fail to form a stable complex (Ishikawa et al., 2017; Mueller et al., 2015; Swamy et al., 2022). Therefore, it is possible that the Kl-Tfc7 protein does not interact properly with other TFIIIC subunits, leading to unstable complex formation, weak target site binding, and subunit degradation.

**Figure 5.**
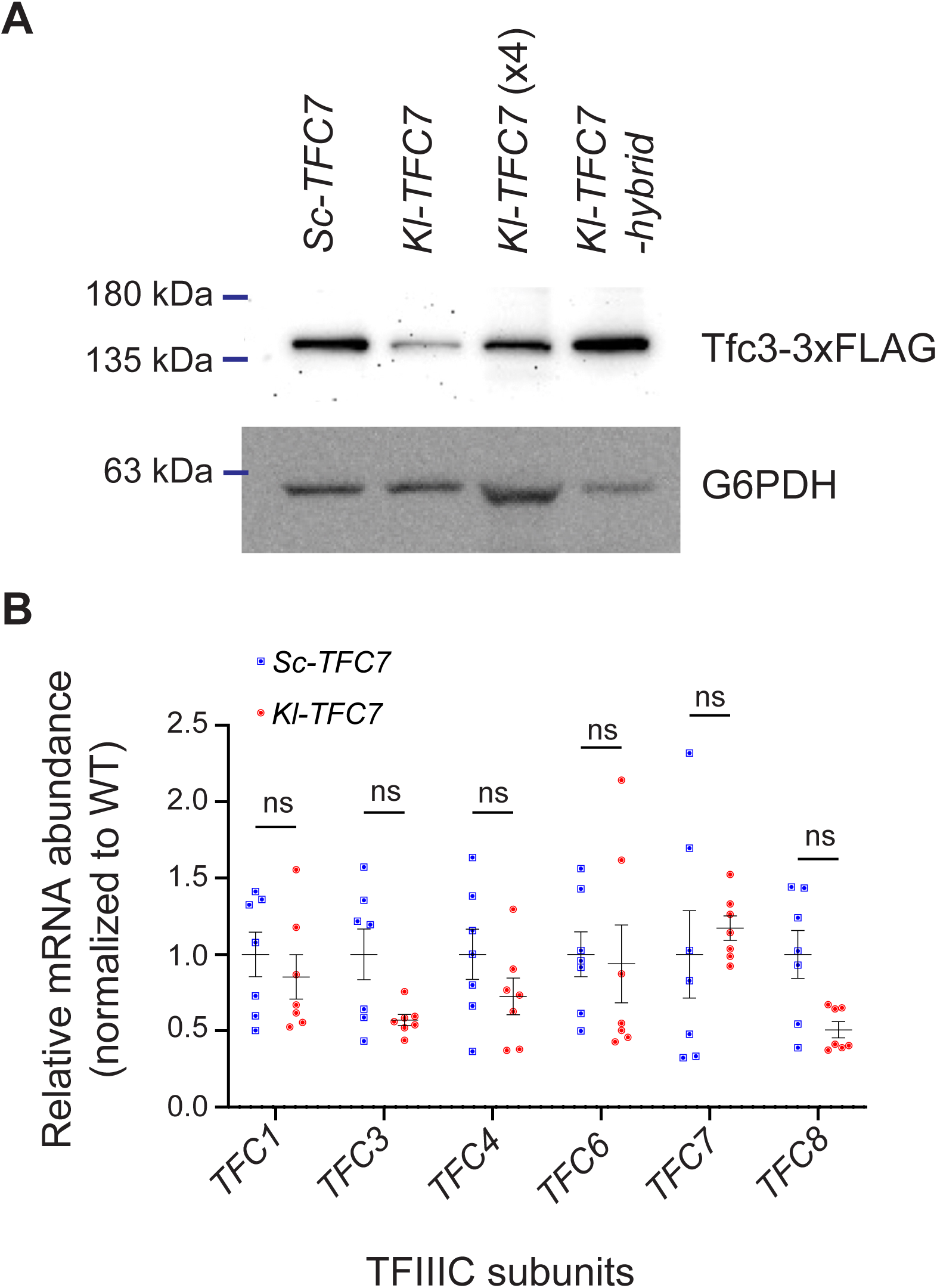
All TFIIIC subunits are expressed normally but Tfc3 protein abundance is reduced in *Kl-TFC7* cells. (A) The Tfc3 protein level in *Kl-TFC7* cells is reduced to about one quarter that of *Sc-TFC7* cells. Total protein lysates were isolated from different strains and hybridized with anti-FLAG and anti-G6PDH antibodies. *Kl-TFC7* (x4) indicates four-fold loading of *Kl-TFC7* lysates. (B) TFIIIC subunits are transcribed to similar levels in both *Sc-TFC7* and *Kl-TFC7* cells. Total RNA was isolated from *Sc-TFC7* and *Kl-TFC7* cells and then the expression of TFIIIC subunits was measured by RT-qPCR. Species-specific primers were used for *TFC7*. The *UBC6* gene was used as an internal control. The mean and standard error are shown. ns, p-value > 0.05 (Student’s t-test, n = 7).

To explore that possibility, first we measured expression of genes encoding the TFIIIC subunits by q-PCR and observed that expression levels were similar between the two strains (Fig. 5B). Thus, introducing *Kl-TFC7* into *S. cerevisiae* cells does not appear to have caused mis-regulation of TFIIIC-linked gene expression. Tfc7 has been shown previously to interact with Tfc1 (Manaud et al., 1998). Recent cryogenic electron microscopy (Cryo-EM) data further indicates that the C-terminal domain of Tfc7 directly interacts with the N-terminal region of Tfc1 to form a triple beta-barrel structure (Supplemental Fig. S5) (Vorländer et al., 2020). A comparison of protein sequences revealed that the C-terminal domains of Sc-Tfc7 and Kl-Tfc7 are highly divergent (Fig. 6A), indicating that this could be the cause of the incompatibility. Indeed, when we used an antibody to pull down Tfc3, Tfc1 co-immunoprecipitated in *Sc-TFC7* cells since all TFIIIC subunits form a stable complex (Fig. 6B). However, Tfc1 could not be pulled down together with Tfc3 in the *Kl-TFC7* background, revealing compromised TFIIIC stability due to the introduction of Kl-Tfc7.

**Figure 6.**
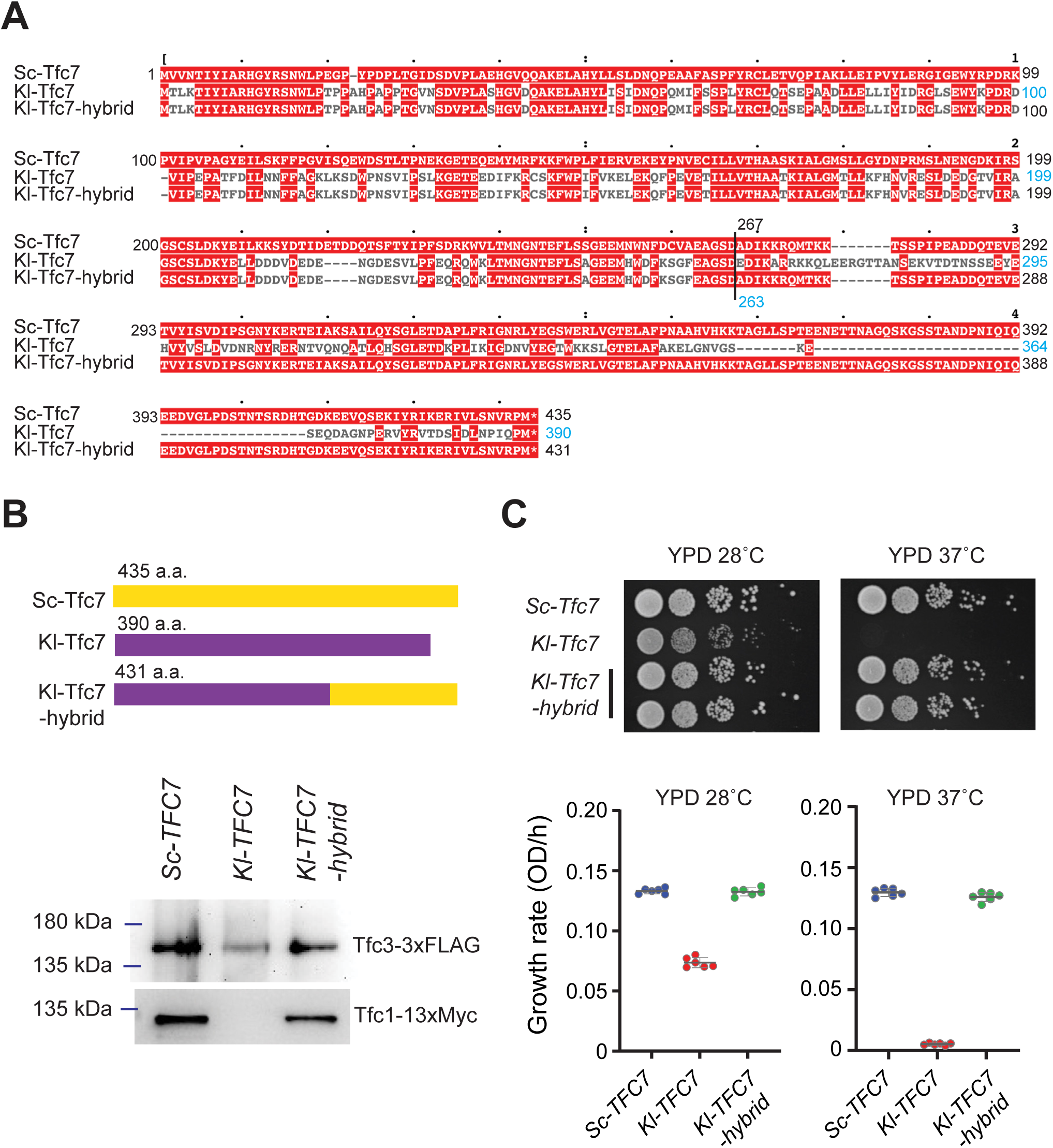
The C-terminal domain of Tfc7 diverges between *S. cerevisiae* and *K. lactis* and causes subunit incompatibility. (A) The C-terminal domain of Kl-Tfc7 diverges significantly from that of Sc-Tfc7. Sequence alignment was performed using ClustalW and identical amino acids are labeled in red. In the Kl-Tfc7-hybrid, the C-terminal domain (a.a. 263-390) of Kl-Tfc7 was replaced by that (a.a. 267-435) of Sc-Tfc7. (B) Swapping the C-terminal domains between Kl-Tfc7 and Sc-Tfc7 rescues the interaction with Sc-Tfc1. Total protein lysates were isolated from different strains. Tfc3-FLAG was immunoprecipitated using anti-FLAG antibody and then the co-immunoprecipiated proteins were hybridized using anti-Myc antibody to detect the interacting Tfc1-Myc protein. (C) Subunit incompatibility is abrogated when the Tfc7-Tfc1 interaction is rescued. Cell fitness was examined using solid and liquid media at 28 °C and 37 °C. *Kl-TFC7-hybrid* cells presented growth similar to *Sc-TFC7* cells in all conditions. The mean and standard error are shown.

To test directly the contribution of the Tfc7-Tfc1 interaction to TFIIIC complex stability, we swapped the C-terminal domain (amino acids 263-390) of Kl-Tfc7 with that of Sc-Tfc7 (a.a. 267-435) to generate a chimeric Tfc7 protein, denoted Kl-Tfc7-hybrid (Fig. 6A). This domain swap resulted in Tfc3 and Tfc1 again co-immunoprecipitating (Fig. 6B), indicating that it had rescued the stability of the TFIIIC complex. Moreover, the Kl-Tfc7-hybrid rescued the protein abundance of Tfc3 (Fig. 5A), supporting the notion that the reduced Tfc3 protein abundance in *Kl-TFC7* cells was due to improper subunit interactions in the TFIIIC complex, followed by protein degradation. *Kl-TFC7-hybrid* cells also exhibited fitness similar to *Sc-TFC7* cells at both normal and high temperatures (Fig. 6C). Together, our data support that the subunit incompatibility is caused by the altered C-terminal domain of Kl-Tfc7, which cannot interact with Sc-Tfc1. Overall then, co-evolution of protein complex subunits allows individual components to accumulate mutations and yet maintain complex stability, which is crucial for target site binding and interactions with other functional components.

## Discussion

In recent years, an increasing number of proteins or complexes have been discovered as possessing multiple functions (Curtis et al., 2023; Singh and Bhalla, 2020). These proteins perform additional functions by interacting with different proteins, DNA, RNA or ligands, and they are involved in various cellular pathways such as the cell cycle, apoptosis, and pathogenesis. Several TFs have also been identified as executing multiple functions (Hall et al., 2004; Levati et al., 2016; Yang et al., 2022). Among them, TFIIIC represents an interesting case since it contains many subunits essential for cell viability and has a very conserved primary function. Previous studies have demonstrated that tRNAs can act as silencing barriers or insulators in various organisms, revealing an involvement of TFIIIC in extra-transcriptional processes (Biswas et al., 2009; Donze and Kamakaka, 2001; Ebersole et al., 2011; Noma et al., 2006; Partridge et al., 2000; Raab et al., 2012; Van Bortle et al., 2014). Genome-wide ChIP-chip experiments have further shown that TFIIIC-binding sites coincide with those of condensin subunits and cohesin loaders, implying a role for TFIIIC in chromosome organization (D’Ambrosio et al., 2008b). Our results reveal that unstable TFIIIC complex formation results in abnormal sister chromatid segregation during mitosis without severely impacting RNA polymerase III transcription. They also demonstrate that these two essential functions of TFIIIC can be uncoupled. Moreover, we have shown that the functional decoupling occurs through the unstable interaction between the C-terminal domain of Kl-Tfc7 and Sc-Tfc1.

When we replaced other *S. cerevisiae* TFIIIC subunits with *K. lactis* orthologs, the cells were not viable, indicating that the whole complex failed to assemble. Partial incompatibility between orthologs provides a useful approach to dissecting the different functions of a protein or complex. Moreover, among the *N. castellii* ortholog replacement lines, we observed that *Nc-TFC1*, *Nc-TFC7* and *Nc-TFC8* were fully compatible with the *S. cerevisiae* background, but that *Nc-TFC4* and *Nc-TFC6* exhibited partial incompatibility. These results provide additional information on the evolutionary trajectories of different subunits, indicating that *Nc-TFC4* and *Nc-TFC6* have undergone more critical changes than the other TFIIIC subunits. This information could not have been revealed by examining the nonsynonymous substitution rate or protein similarity score (Supplementary Table S1).

The role of TFIIIC in facilitating RNA polymerase III transcription is conserved from yeast to humans. However, all six TFIIIC subunits proved partially or fully incompatible between *K. lactis* and *S. cerevisiae*. Thus, many changes have accumulated within this complex and yet its primary function has been maintained. How has that been accomplished? One possibility is that the microenvironment provided by multi-subunit complexes allows the subunits to change their interaction interfaces more quickly. A previous non-biased screen identified several rapidly-evolving essential genes that all encode subunits of large protein complexes (Lai et al., 2023). In that study, it was hypothesized that when a subunit interacts with multiple components of the complex, it creates a robust microenvironment. Individual interface mutations can be tolerated (without compromising fitness) if stability of the entire complex is maintained by other interactions. Such changes may gradually lead to co-evolution of interacting subunits, resulting in conformational changes without losing the original function. Nevertheless, the altered conformation may open up the possibility of incorporating new subunits, thereby increasing complexity or/and evolving novel functions. This hypothetical scenario was supported by experiments on the anaphase-promoting complex/cyclosome (APC/C) (Lai et al., 2023). In the case of TFIIIC, we have found that swapping the C-terminal domain of Tfc1 rescued subunit incompatibility, but replacing *TFC1* and *TFC7* with *K. lactis* orthologs simultaneously failed to do so (Supplementary Table S1). Therefore, the interactions between Tfc1 and non-Tfc7 subunits are also likely crucial for the stability or function of the entire complex. The complex microenvironment hypothesis thus represents an attractive model for how a protein complex enhances its moonlighting potential and can evolve multiple functions.

Our data have demonstrated that essential complex subunits may change their interaction interfaces after species have diverged. The initial changes likely result from genetic drift. However, selection may be involved in accelerating the evolutionary change once a new function or regulation has emerged. The experiments presented herein do not test if the conformation of TFIIIC has changed while preserving its original functions or if in different species it has evolved differential regulation or novel functions. To address that question, we would need to directly examine and compare its components, dynamics and functions between different species. Nonetheless, our results demonstrate that essential proteins or complexes are not static units and can diverge to evolve additional functions even if their primary functions are conserved among species. The multi-functionality of a protein may play a crucial role in driving its evolution.

## Acknowledgments

We thank Dr. Hung-Ta Chen, Dr. Rey-Huei Chen, and Dr. Cheng-Fu Kao for yeast strains, plasmids, and technical assistance. We thank members of the Leu lab for helpful discussions and comments on the manuscript. We thank the Genomics, Bioinformatics, and Imaging Cores of IMB for technical assistance, and John O’Brien for manuscript editing. JYL was supported by Academia Sinica of Taiwan (grant no. AS-IA-110-L01 and AS-GCS-113-L03) and the National Science and Technology Council of Taiwan (NSTC 113-2326-B-001-002).

## Author contributions

JYL conceived the study. AG and JYL designed analyses and interpreted results. AG, PCH, and RRRL performed the experiments. AG and JYL wrote the paper.

## Competing interests

The authors have declared that no competing interests exist.

## Supplementary figure legends

**Supplementary Figure S1.**
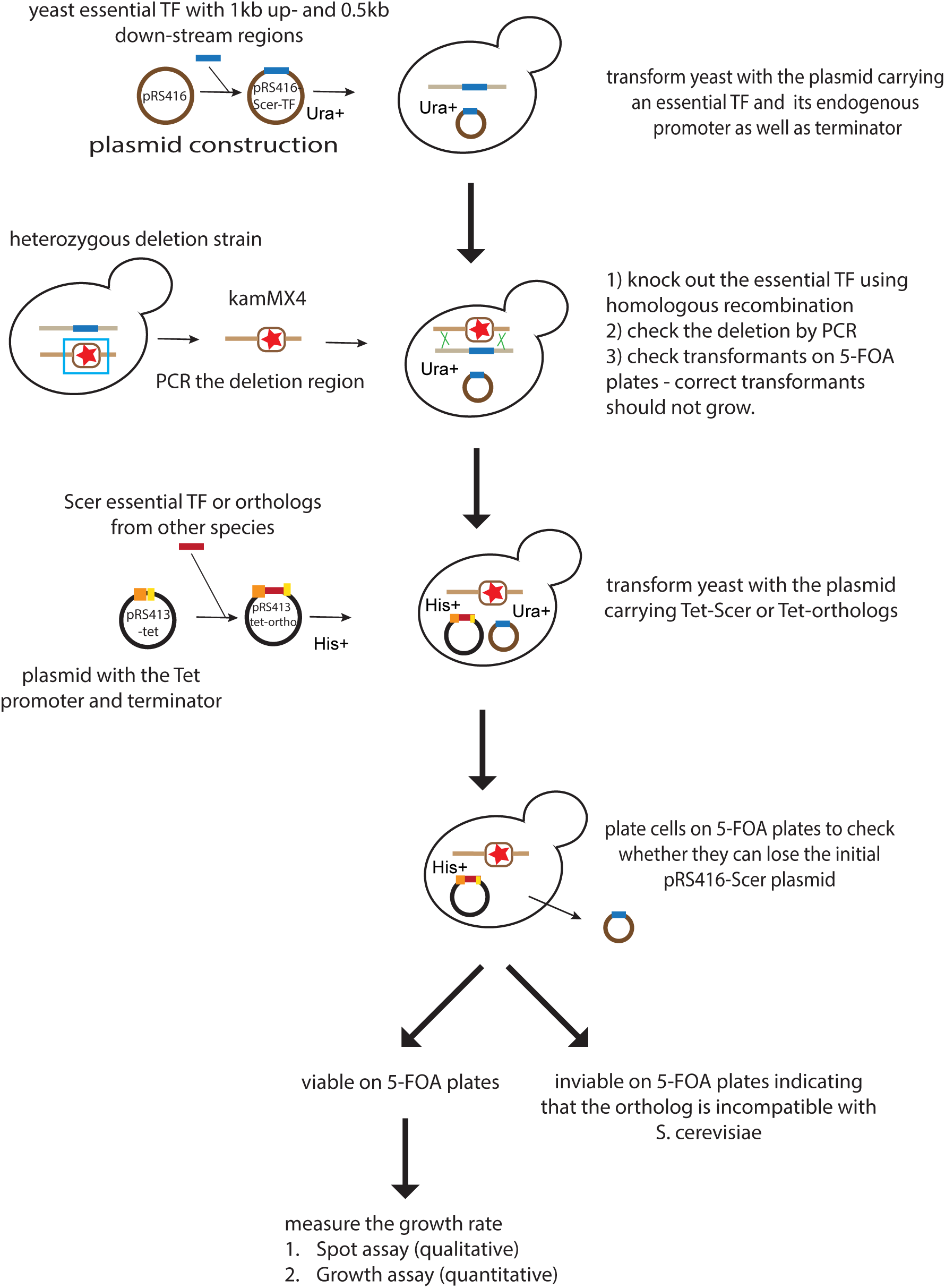
Experimental protocol for generating eTF replacement lines. First, we constructed the plasmid containing the target eTF (pRS416-*Sc-eTF*) and transformed it into the shuffling strain. The endogenous *S*. *cerevisiae* eTF was then knocked out by means of homologous recombination. Next, we built the orthologous gene-containing plasmids (pRS413-pTetO_7_-*Nc-eTF* or pRS413-pTetO_7_-*Kl-eTF*) and used them to replace the pRS416-*Sc-eTF* plasmid. If the essential gene could be replaced by its orthologs, the growth rates of the replacement lines were measured.

**Supplementary Figure S2.**
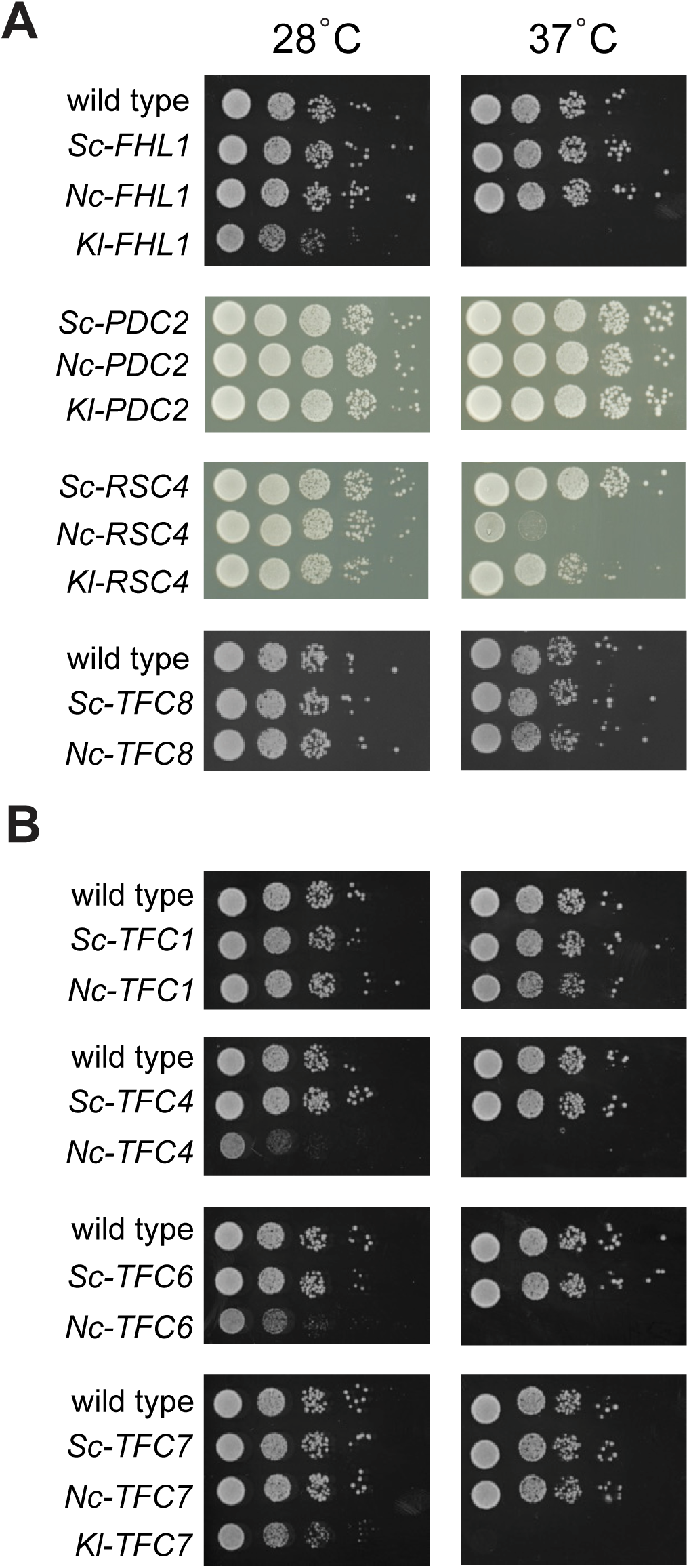
Spot assay data for selected replacement lines. (A) *S*. *cerevisiae FHL1*, *RSC4*, *PDC2*, and *TFC8* genes were replaced by their orthologs derived from *N. castellii* or *K. lactis*. The viable replacement lines were grown on rich medium plates (YPD) at 28 °C (optimal temperature) and 37 °C (heat stress). (B) *S*. *cerevisiae* TFIIIC subunit-encoding genes, i.e., *TFC1*, *TFC4*, *TFC6*, and *TFC7*, were replaced by their orthologs from *N. castellii* or *K. lactis*. The viable replacement lines were grown on YPD at 28 °C and 37 °C.

**Supplementary Figure S3.**
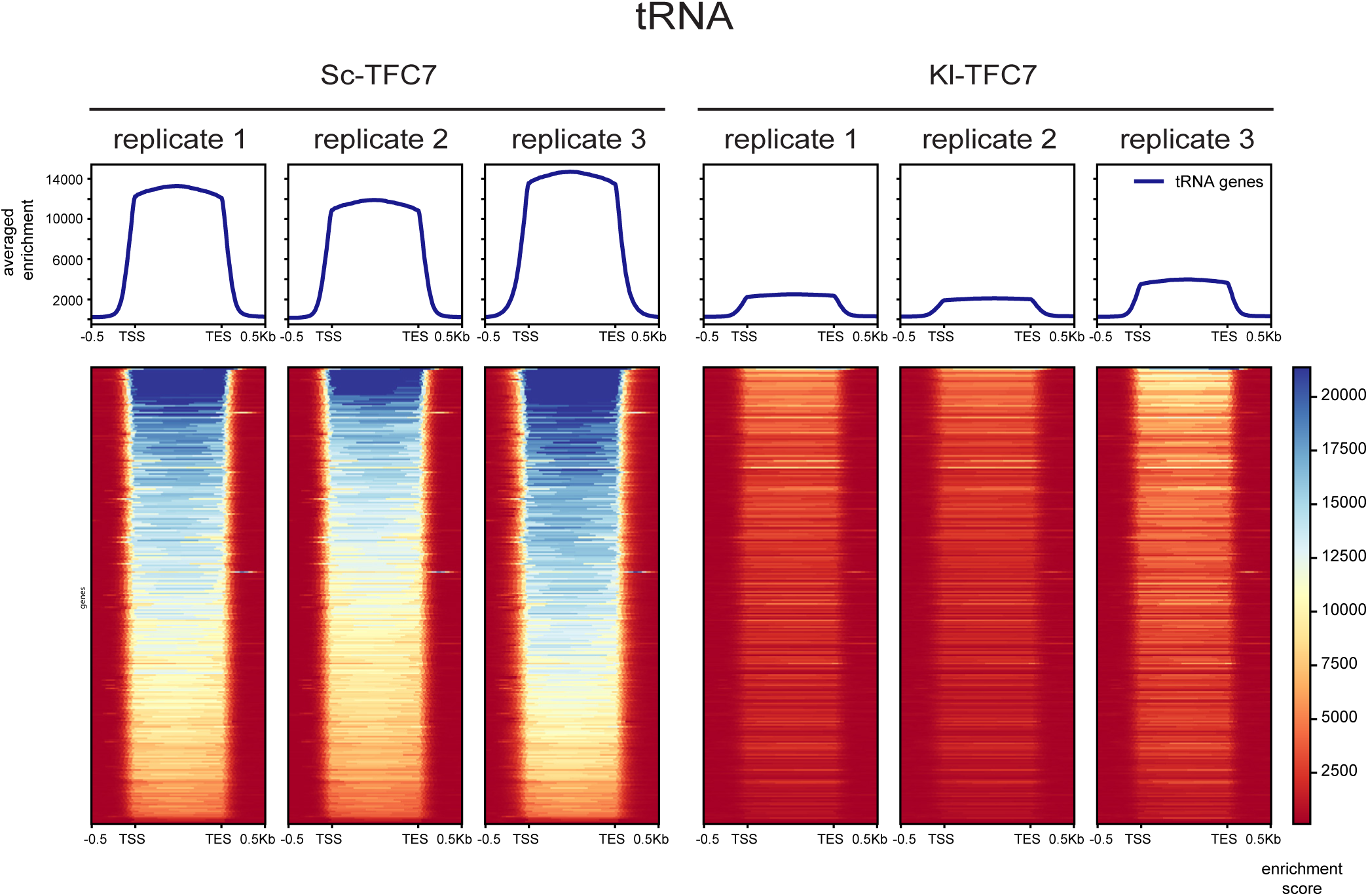
Binding of Tfc3 to tRNAs is significantly reduced in *Kl-TFC7* cells. The top panel represents an average of enriched peaks at all tRNA genes for three biological replicates of the *Sc-TFC7* and *Kl-TFC7* lines. TSS indicates the beginning of the tRNA gene and TES indicates 1000 bp downstream. The heatmap represents all individual tRNA peaks (n = 275).

**Supplementary Figure S4.**
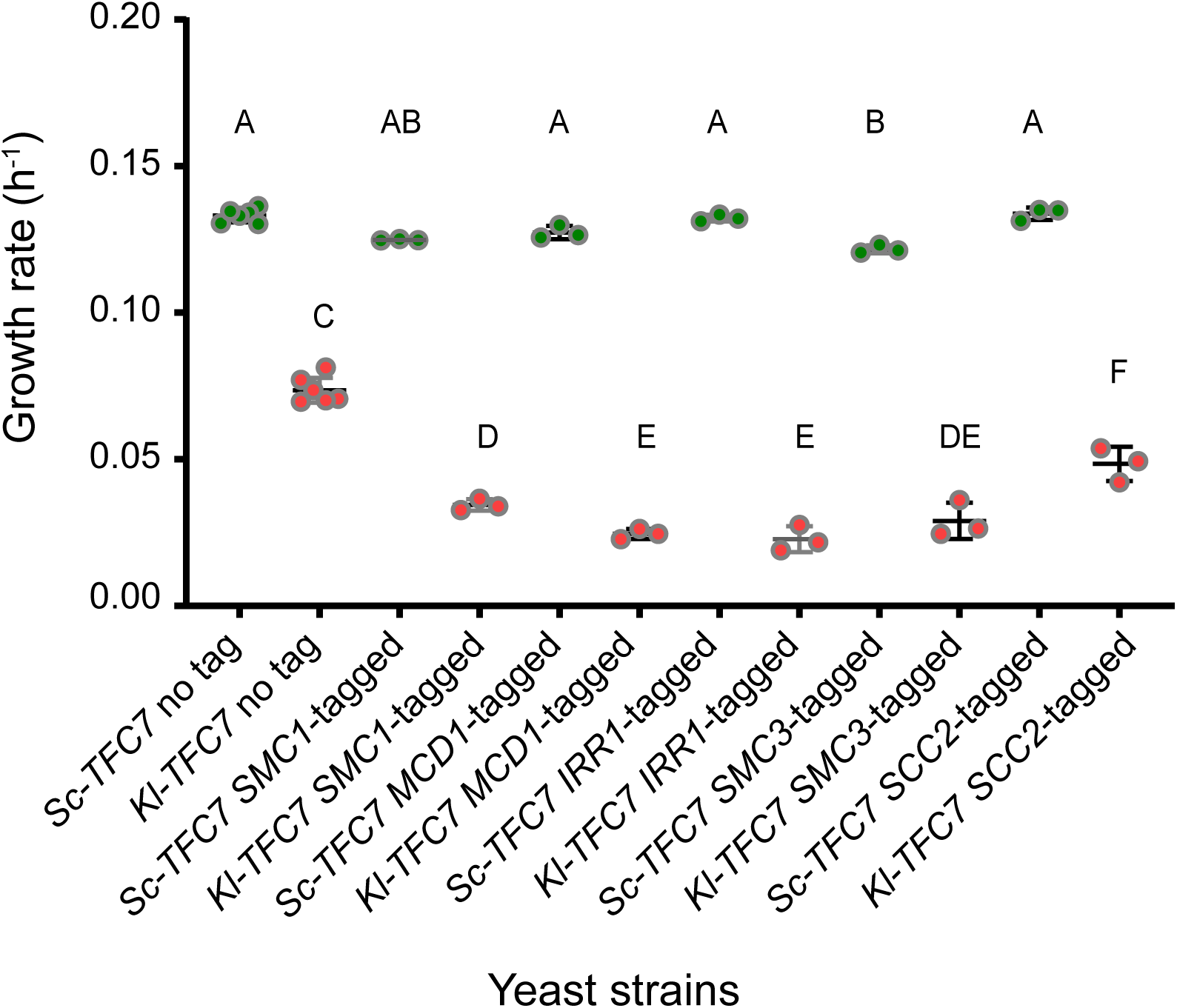
Tagged cohesin subunits and loading factors cause severe synthetic sickness in the *Kl-TFC7* background but not in the *Sc-TFC7* background. Growth rate at 28 °C in YPD (rich medium) for replacement lines with or without additional tags. Cohesin proteins were tagged with the TAP epitope (Smc1, Mcd1, Irr1 and Smc3) or 13Myc epitope (Scc2). Cells were grown in YPD at 28 °C and their growth rates were measured by plate readers. The mean and standard error are shown. Different letters above each sample indicate statistically significant differences (ANOVA and Tukey’s HSD post-hoc test, adjusted p-value < 0.05).

**Supplementary Figure S5.**
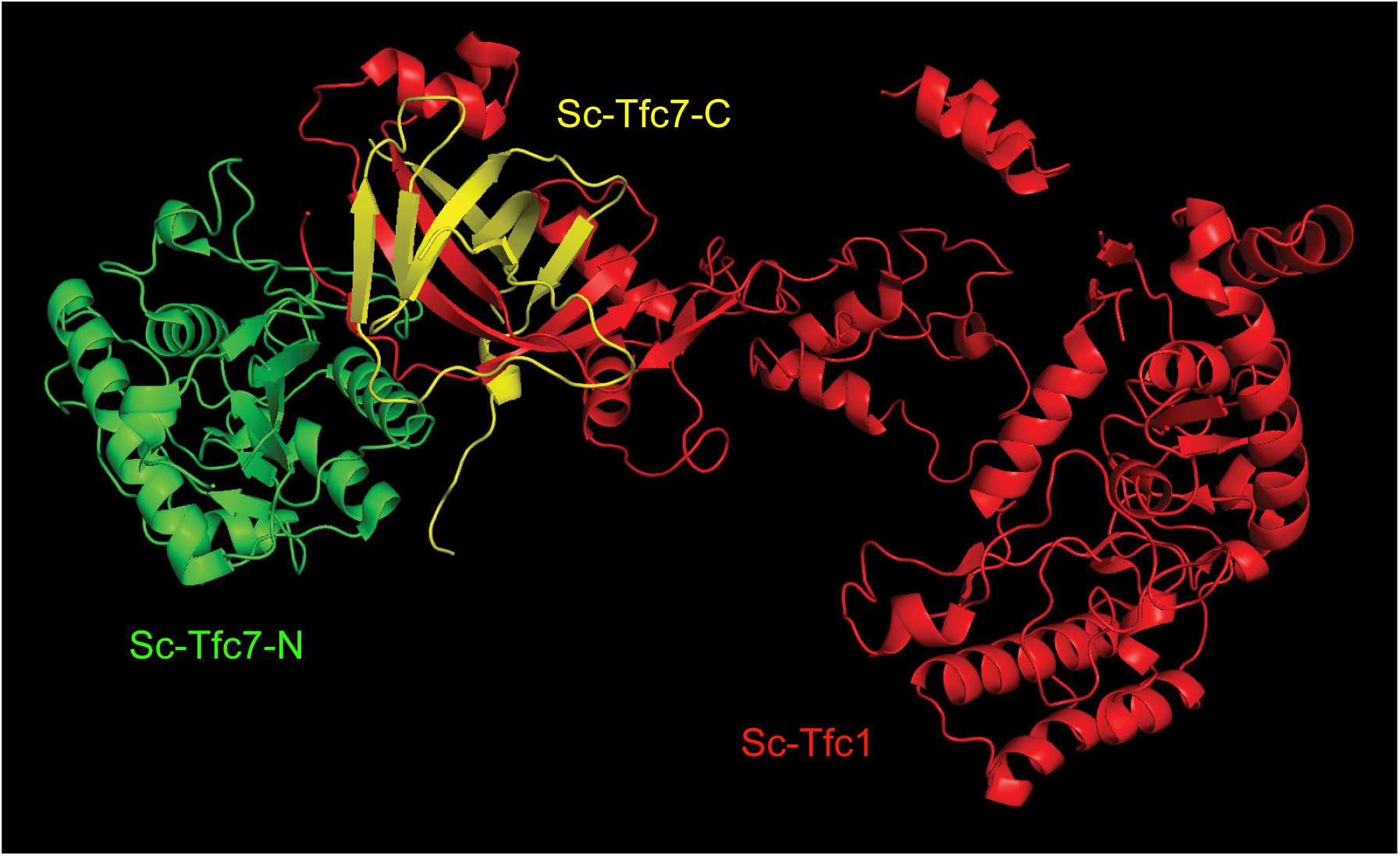
The interaction between Tfc7 and Tfc2. The structure of the TFIIIC subcomplex τA 6yj6 consisting of Sc-Tfc4, Sc-Tfc1 and Sc-Tfc7, as determined by cryo-EM to a resolution of 3.1Å, was provided by Vorländer Matthias K. (https://doi.org/10.1038/s41467-020-18707-y). Sc-Tfc4 has been removed from the figure to increase clarity, with Sc-Tfc7 divided into two colors to represent its N-terminal (green) and C-terminal (yellow) domains. The C-terminal domain of Sc-Tfc7 adopts a beta-barrel structure that forms a τ95–τ55 dimer with Sc-Tfc1.

## Materials and Methods

### Strain and plasmid constructions

All the *S*. *cerevisiae* strains were derived from R1158 (*MATa met15-0 ura3::CMV-tTA-ura3 his3-1 leu2-0 trp1-63*) (Mnaimneh et al., 2004). Yeast cells were normally grown at 28 °C in yeast extract-peptone-dextrose (YPD: 1% yeast extract, 2% bacto peptone, and 2% glucose) or dextrose synthetic dropout media if not specifically mentioned otherwise. Gene deletions and tagging were done using standard polymerase chain reaction (PCR)-based recombination methods, followed by confirmation by PCR and sequencing.

To construct the ortholog replacement lines, a pRS416 plasmid with an essential transcription factor (pRS416-*Sc-eTF*) was transformed into the *S. cerevisiae* R1158 strain and the corresponding eTF was deleted from the genome (Supplementary Figure 1). Then, we transformed the strain with the pRS413-pTetO_7_-ortholog plasmids and selected the transformants on 5-Fluoroorotic acid (5-FOA) plates to obtain the ortholog-replacement strains. All ortholog-replacement strains were checked using PCR to confirm loss of pRS416-*Sc-eTF*. If no colony grew on the 5-FOA plates after multiple trials, we assumed that the orthologous gene was completely incompatible with the host. Multiple clones were selected and maintained for each step. The orthologous gene information of *N. castellii* and *K. lactis* was obtained from the Yeast Gene Order Browser (http://ygob.ucd.ie/) (Byrne and Wolfe, 2005).

To construct the strains for monitoring chromosome segregation, a pRS405 plasmid containing tetR-GFP (Chen, 2019) was linearized with *Afl*II and integrated into the *LEU2* locus. Next, a pRS406 plasmid with a tetOx112 array was digested with *Afl*II and integrated into the YDR246W-A locus (Chen, 2019). After the integrations were confirmed by PCR, the strains were used for GFP-dot assays.

To construct the *Kl-TFC7*-*hybrid* gene, we swapped the C-terminal domain of Kl-Tfc7 with that of Sc-Tfc7. We amplified by PCR the N-terminal fragment of *Kl-TFC7* with an overhang, as well as the C-terminal fragment of *Sc-TFC7* with an overlapping overhang to that of *Kl-TFC7*. PCR fragments extracted from the gel were ligated with linearized *pRS413-*pTetO_7_ plasmid DNA using In-Fusion cloning. Transformed colonies were checked using PCR and Sanger sequencing. Confirmed plasmids were transformed into the shuffle strain to replace the *pRS416*-*Sc-TFC7* plasmid.

### Growth rate measurement

For spot assays, cells grown in YPD overnight at 28 °C were harvested by centrifugation and serially diluted to the desired cell densities with sterile ddH_2_O. Five μl of each dilution (10^7^–10^3^ CFU/ml) was spotted onto the agar plates and incubated at 28 °C or 37 °C. The plates were scanned every day for 5 days and the image records were kept.

To measure the growth rate in liquid medium, cells grown in YPD overnight at 28 °C were inoculated into 120 μl YPD medium in a 96-well plate (Tissue Culture Testplate 96 wells F-bottom, #92096, TPP, Trasadingen, Switzerland) at a cell density of 0.2 OD_600_/ml. Cell growth was measured at OD_595_ in “2x2 multiple reads per well” mode every 12 min using a plate reader (Infinite 200 PRO, Tecan, Mannedorf, Switzerland). The Tecan software Magellan Version 7.2 was used for data acquisition and analyses. In Magellan, we selected the plate definition “[TPP96ft]-Tecan Plastic Products AG6 Flat Transparent”. Each 12-min cycle included 3 min reading, 1 min shaking, 3 min standing, 1 min shaking, 3 min standing, and 1 min shaking.

### Total RNA isolation

Total RNA was extracted according to the phenol-chloroform method, as previously described with some modifications (Hsu et al., 2021). In brief, early log-phase (0.5–0.8 OD_600_/ml) cells from subcultures grown in YPD were harvested by centrifugation, washed once with ddH_2_O, and stored at −80 °C. Approximately 10 OD_600_ of cells were thawed on ice and resuspended in 500 μl of nuclease-free and ice-cold lysis buffer (0.1 M Tris-Cl [pH 7.5], 0.1 M LiCl, 2% beta-ME, 0.01 M EDTA, and 5% SDS). The mixtures were transferred to a microcentrifuge tube containing ice-cold 500 μl PCIA (phenol:chloroform:isoamyl alcohol = 25:24:1, pH 4.5; 0.1% 8-Hydroxyquinoline) and 0.5 g glass beads (#11079105, BioSpec Products, Bartlesville, USA). The cells were lysed by vortexing them at the maximum speed for 5 min. Supernatants were obtained by centrifugation for 10 min at 4 °C. Each aqueous phase was extracted by two more rounds with 200 μl ice-cold PCIA followed by mixing for 3 min and centrifugation. Each final aqueous phase (350–400 μl) was transferred into 1 ml of 99% ethanol for precipitation with mild mixing for 2 h. Nucleic acid pellets were spun down, washed with 70% ethanol, air-dried for 5 min, and then resuspended in 50 μl nuclease-free water (UltraPure™ DNase/RNase-Free Distilled Water, #10977015, Invitrogen by Thermo Fisher Scientific, Waltham, USA).

DNA contaminants in the total RNA were removed using a TURBO DNA-free^TM^ kit (#AM1907, Invitrogen by Thermo Fisher Scientific) by treating 40 μg of total RNA at 37 °C for 40 min with 2 μl of TURBO DNase in a 50-μl reaction. The DNase was removed by treating the reaction mixture with 5 μl of inactivation reagent beads for 15 min at room temperature with mild vortexing every 5 min. The purified RNA in the supernatants was collected by spinning down the beads and quantified by measuring the OD_260_ using an ND-1000 spectrophotometer (Thermo Fisher Scientific). Final RNA quality was checked using a Bioanalyzer 2100 instrument (Agilent Technologies, Santa Clara, USA) with an RNA 6000 Nano LabChip kit (Agilent Technologies).

### Nascent RNA isolation

For nascent RNA isolation, cells were cultivated in 100 ml YPD at 28 °C until the OD_600_ reached ∼0.8. The newly synthesized RNAs were labeled for 6 min by adding freshly prepared 4-thiouracil (Sigma-Aldrich) to a final concentration of 5 mM. Concurrently, *S. pombe* cells were cultivated under analogous conditions in YES medium at 31 °C and subjected to the same labeling process to serve as a spike-in control across all experimental samples. After labeling, the cells were pelleted, flash-frozen in liquid nitrogen, and subsequently stored at −80 °C for future applications.

Prior to total RNA extraction, *S. cerevisiae* and *S. pombe* cells were combined in a ratio of 3:1. Total RNA was extracted according to a phenol-chloroform method described previously (Hsu et al., 2021). Before the process of biotinylation, the RNA samples were heated for 10 min at 60 °C and subsequently cooled on ice for an additional 5 min. A total of 200 μg of RNA underwent biotinylation utilizing 200 μl of 1 mg/mL EZ-link HPDP-Biotin (Pierce) in conjunction with 100 μl of biotinylation buffer (100 mM Tris-HCl pH 7.5, 10 mM EDTA), with the solution being adjusted to a final volume of 1000 μl using DEPC-treated RNase-free water (Sigma-Aldrich) for a duration of 3 hours at ambient temperature. Following chloroform extraction and isopropanol precipitation (1/10 volume 5M NaCl and 2.5 volumes isopropanol), the purified RNA was reconstituted in 100 μl of DEPC-treated RNase-free water (Sigma-Aldrich).

Recovered RNA samples were incubated at 65 °C for 10 min and cooled on ice for 5 min. Biotinylated RNAs were then bound to μMACS streptavidin microbeads (Miltenyi Biotec, Bergisch Gladbach, Germany) for 90 min at room temperature. Purification was performed using a μMACS streptavidin starting kit (Miltenyi Biotec), with columns equilibrated with a washing buffer. Samples were passed through the columns twice and washed five times with increasing volumes of washing buffer. Labeled RNAs were eluted twice with 200 μl of 100 mM DTT. After overnight ethanol precipitation, RNAs were washed in 70% ethanol and resuspended in DEPC-treated RNase-free water (Sigma-Aldrich). Samples were stored at −80 °C for future use (Baptista et al., 2017).

### RNA gel

Total tRNA levels were visualized using a 12% TBE urea gel (5X TBE, 12% Urea, 30% Acryl/Bis (19:1), 10% ammonium persulphate solution (APS), TEMED). To warm up the gel, it was pre-run in 0.5X TBE buffer for 30 min, and excess urea collected in the wells was removed by thorough pipetting. Then, 2-4 μg of sample was loaded in each well, along with RNA Gel Loading Dye (2X) (# R0641, Thermo Fisher Scientific). The gel was run at a constant 100V for 5 h, followed by EtBr staining in a staining tank for 10 min. Single bands for 5.8S ribosomal RNA were observed between the 200-300 bp marker, followed by 5S ribosomal RNA and three bands of tRNA at the bottom of the gel, i.e., above and below the 100 bp marker. The intensity of the total tRNA band was normalized with 5.8S rRNA because 5.8S rRNA is transcribed by RNA polymerase I and should not be influenced by the ortholog replacement of TFIIIC subunits.

### Reverse transcription (RT) and quantitative PCR (q-PCR) analyses

To measure tRNA levels, 1 μg of RNA was reverse-transcribed using a RevertAid First Strand cDNA Synthesis Kit (#K1621, Thermo Fisher Scientific). To minimize the effect of secondary structure, the reaction was conducted at 60 °C (instead of 45 °C) and extended for 30 min (instead of 15 min) to reduce the effect of transcription pauses due to tRNA modifications. For nascent tRNAs, cDNA synthesis was performed on 10 μl of labeled RNA using random hexamers and High-Capacity cDNA Reverse Transcription Kits (Thermo Fisher Scientific).

Real-time qPCR was performed with the model 7500 Fast Real-Time PCR System and the model QuantStudio^TM^ 12K Flex Real-Time PCR System (Applied Biosystems by Thermo Fisher Scientific). In brief, each 10-μl reaction mixture contained 100 ng cDNA, 150 nM (each) primers, and 10 μl Fast SYBR^TM^ Green Master Mix (#4385612, Applied Biosystems by Thermo Fisher Scientific). The reactions were performed with one cycle at 95 °C for 10 min, followed by 45 repeated cycles at 95 °C for 15 s, and 60 °C for 1 min. The *RDN58* transcripts encoding 5.8S rRNA were used as an internal control in the q-PCR of steady-state tRNA experiments and *S. pombe* tubulin transcripts were used as a spike-in control in the q-PCR of nascent tRNA experiments. The average ΔΔC_T_ and standard deviation were determined from at least three biological replicates, each containing three technical repeats. The relative fold-change of each gene is shown according to the 2^−CT^ method.

To measure the abundance of TFIIIC complex mRNA, 1 μg of RNA was reverse-transcribed using a High-Capacity cDNA Reverse Transcription Kit (#4368814, Thermo Fisher Scientific). The *UBC6* transcripts were used as an internal control for the q-PCR. The average ΔΔC_T_ and standard deviation were determined from at least three biological replicates, each containing three technical repeats. The relative fold-change of each gene is shown according to the 2^−CT^ method.

For ChIP-qPCR, extracted samples called Input-DNA and ChIP-DNA were diluted to 0.05 ng and 0.5 ng ratios to be used as templates. Primers were designed using Primer3 and Oligo Calc (http://biotools.nubic.northwestern.edu/OligoCalc.html) with the Forward primer within the tRNA gene and the Reverse primer outside the gene to ensure a single primer binding site. All primers used in the q-PCR experiments are listed in Supplementary Table S3.

### Chromosome segregation assays

For GFP-dot assay, yeast cells were cultured overnight in 2x complete supplement medium (2x CSM: 0.7% yeast nitrogen base without amino acids, 2% dextrose, 0.2% CSM powder, pH 7) and refreshed for 4 h before observation. Multiple clones were generated for both the *Sc-TFC7* and *Kl-TFC7* strains, in which a tetO array of 112 repeats was integrated into chromosome IV. When tetR-GFP binds to the tetO array, it emits a sharp green fluorescence dot, which can be detected under fluorescence microscopy. Each clone for both *Sc-TFC7* and *Kl-TFC7* strains was imaged on different days, and imaging data was collected for further analysis using ImageJ.

### Chromatin immunoprecipitation

*Kl-TFC7* and *Sc-TFC7* cells with the tagged Tfc3 protein (Tfc3-3XFlag-6XHis) were grown in the YPD medium at 28 °C overnight and subsequently diluted in YPD to a cell density of 0.2 OD_600_/ml. After incubation at 28 °C, mid log-phase cells (0.5 OD_600_/ml in YPD) were fixed with 1% formaldehyde at 25 °C for 10 min with shaking at 120 rpm and then quenched with 125 mM glycine at 25 °C for 5 min with shaking at 120 rpm. All subsequent steps were performed in an ice-cold or 4 °C environment. Cells were harvested, washed twice with TBS (20 mM Tris-Cl, pH 7.5; 150 mM NaCl), and stored at −80 °C until use.

To break cells, 500 OD_600_ of cells were thawed and resuspended in 4 ml of FA buffer (50 mM HEPES, pH 7.5; 140 mM NaCl; 0.1% SDS; 1 mM EDTA; 1% Triton X-100; 0.1% Deoxycholate, Na salt) with 1/100 PIC (Protease inhibitor cocktail Set IV, in DMSO; #539136, Merck, Darmstadt, Germany) and 1 mM phenylmethanesulfonyl fluoride (PMSF, #P7626, Sigma-Aldrich, St. Louis, USA). The cell suspension was divided into four 2-ml breaking tubes (BioSpec products, #10832, Microtube 2 ml with cap) containing a 1-ml volume of glass beads (#11079105, BioSpec Products). Lysis was performed by eight cycles of 1-min beating, followed by 1-min chilling on ice (BioSpec Mini-BeadBeater-16, BioSpec Products). Cell lysates and glass beads were separated by punching a hole in the bottom of the tube with a red-hot 18G needle and then performing slow centrifugation (500 x *g*, 3 min, 4 °C; Eppendorf 5810R centrifuge, A-4-62 rotor). The collected lysates were combined into a 15-ml centrifuge tube and washed with two cycles of 5 ml FA buffer, followed by centrifugation (12K rpm, 5 min, 4 °C; Eppendorf 5810R centrifuge, F-34-6-38 rotor). The washed lysates were completely resuspended in 2 ml FA buffer with 1/100 PIC and 1 mM PMSF in a new 15-ml centrifuge tube (#CFT011150, JET BIOFIL, Guangzhou, China). Chromatin was sheared by sonication in a Bioruptor water bath sonicator (Diagenode, Liege, Belgium) with 15 ml tube-chip units (high intensity; 30 sec on, 30 sec off, 15 min/cycle; total three cycles) at 4 °C. The sheared lysate was centrifuged at 12000 x *g* for 10 min at 4 °C (Eppendorf 5810R centrifuge, F-34-6-38 rotor). Approximately 25 μl of supernatant corresponding to 5 OD_600_ of cells was collected, mixed with 200 μl TES buffer (50 mM Tris-Cl, pH 8; 10 mM EDTA, pH 8; 1% SDS) plus 175 μl TE buffer (10 mM Tris-Cl; 1 mM EDTA, pH 8), and stored at -20 °C as the input DNA.

Immunoprecipitation was performed on *Sc-TFC7* and *Kl-TFC7* replacement line cells with Tfc3-3XFlag-6XHis by incubating all the remaining sheared chromatin with 60 μl Anti-FLAG^®^ M2 Magnetic Beads (#M8823, affinity-isolated antibody, Merck) in 6 ml of FA buffer with 1/100 PIC and 1 mM PMSF. Binding was performed in a 15-ml tube with mixing using an end-over-end rotator (Intelli-Mixer, RM-2L, F1 mode, 30 rpm) at 4 °C for 16 h. The magnetic beads were anchored with a DynaMag^TM^-2 Magnet and the bound complexes were washed four times for 5 min with 1 ml of each wash buffer [FA buffer × 1, high-salt FA buffer (50 mM HEPES, pH 7.5; 500 mM NaCl; 0.1% SDS; 1 mM EDTA; 1% Triton X-100; 0.1% Deoxycholate, Na salt) × 1, DOC buffer (10 mM Tris·Cl; 1 mM EDTA, pH 8; 25 0mM LiCl; 0.5% IGEPAL^®^ CA-630; 0.5% Deoxycholate, Na salt) × 1, and TE buffer × 1] with mixing (Intelli-Mixer, RM-2L, F1 mode, 30 rpm) at room temperature. The bound complexes were eluted twice by heating at 65 °C in 200 μl TES buffer for 20 min and in 200 μl TE buffer for 10 min. Two eluates were pooled as the ChIP-DNA. To remove RNA, both the input DNA and ChIP-DNA (∼400 μl each) were treated with 5 μl of 10 mg/ml RNase A (R5503, Sigma; 100 mM Tris-Cl, pH 7.4) at 37 °C for 30 min. For de-crosslinking, both the RNase-treated input DNA and ChIP-DNA were treated with 40 μl of 20 mg/ml Proteinase K (#1.24568.0500, Merck; in 50 mM Tris-Cl, pH 8) at 42 °C for 1 h and then at 65 °C for 16 h. De-crosslinked DNA was purified using the QIAquick DNA Purification Kit (#28106, Qiagen, Hilden, Germany) according to the manufacturer’s instructions, with a modification of a two-cycle wash step using the PE buffer. Final ChIP-DNA and input DNA were eluted with 50 μl EB buffer. DNA concentrations were measured by using a Qubit™ dsDNA HS Assay Kit (#Q32854, Invitrogen by Thermo Fisher Scientific) (Hsu et al., 2021).

### ChIP-sequencing analyses

ChIP-seq analysis was performed from three biological replicates of YPD-grown cells. The average fragment length of sonicated fragments was 150–1000 bp. For each condition, libraries were prepared from 3.96–20 ng of ChIP-DNA and input-DNA using the KAPA LTP Library Preparation Kit for Illumina^®^ platforms (#KK8232, KAPA Biosystems by Roche, MA, USA) according to the manufacturer’s instructions. Notably, after adapter ligation, double-sided size selection between 250–450 bp for adapter-ligated fragments was performed using KAPA Pure Beads (#KK8000/07983271001, KAPA Biosystems by Roche). Size-selected fragments were then PCR-amplified for 12 cycles. The size of each library was assessed using a Bioanalyzer 2100 instrument (Agilent Technologies) with a High Sensitivity DNA Kit (Agilent Technologies). The concentration of each library was quantified by using a Qubit™ dsDNA HS Assay Kit (#Q32854, Invitrogen by Thermo Fisher Scientific) and q-PCR. Single-read sequencing (75 bp) of the libraries was performed using a NextSeq 500/550 high output reagent kit V2_75 cycle (#FC-404-2005, Illumina, San Diego, USA) on an Illumina NextSeq500 sequencer.

Quality control and adaptor trimming of sequencing reads were processed in FastQC version 0.11.8 (https://www.bioinformatics.babraham.ac.uk/projects/fastqc/). Reads were mapped to the *S. cerevisiae* R64-3-1 genome using Bowtie2 (Langmead and Salzberg, 2012). To determine which genomic regions are enriched for Tfc3 binding, Bowtie2 mapped BAM files were detected for the ChIP peaks using MACS (Zhang et al., 2008). Only peaks detected in all three biological repeats with p < 0.05 were used for the downstream analysis.

For the peak-to-gene assignments, the peaks were assigned to specific genes based on the location of each peak center. Only the peaks with centers located within the same region as detected peaks were assigned. The gtf and bed files containing all areas in the genome were used to assign non-coding RNA, tRNA, ARS, TyLTR and centromere. These files were downloaded from SGD (https://www.yeastgenome.org/).

A Java tool developed by Babraham Bioinformatics (https://www.bioinformatics.babraham.ac.uk/publications.html) called SeqMonk was used to create the tables needed to make the scatter plot. For the supplementary figure, Deeptools was used to make the tRNA matrix and heatmaps (Ramirez et al., 2016).

### Total protein extraction

Total yeast protein extraction was conducted according to the TCA protocol. The yeast cells were collected and centrifuged to remove supernatant. The cell pellets were then resuspended in ice-cold sterile ddH_2_O and washed again. After discarding the supernatant, the pellets were resuspended in lysis buffer (0.185 M NaOH, freshly added 0.75% β**-**mercaptoethanol) and incubated on ice for 10 min. Then, 150 μl of 55% TCA solution was added for protein precipitation via an additional incubation on ice for 10 minutes, followed by centrifugation to collect the protein pellets. The pellets were dissolved in 44.5 μl HU sample buffer (8M Urea; 5% SDS; 0.2M Tris-Cl, pH 6.5; 1 mM EDTA, pH 8.0; 0.01% Bromophenol blue) plus 2.5 μl β**-**mercaptoethanol plus 3 μl 2 M Tris base, making a total of 50 μl per OD of cells, which was heated at 65 °C for 40 min with mixing in between, and centrifuged before loading onto a polyacrylamide gel for analysis.

### Protein co-immunoprecipitation

*Kl-TFC7* and *Sc-TFC7* cells were grown overnight in YPD medium at 28 °C and subsequently diluted in YPD to a cell density of 0.2 OD_600_/ml. After incubation at 28 °C with shaking at 120 rpm, cultures of mid-log phase cells (0.7-0.8 OD_600_/ml in YPD) were harvested. Harvested cells were washed twice with ice-cold TBS (20 mM Tris-Cl, pH 7.5; 150 mM NaCl), and stored at −80 °C until use.

To break cells, 500 OD_600_ of cells were thawed and resuspended in 3 ml of FA buffer (50 mM HEPES, pH 7.5; 140 mM NaCl; 0.1% SDS; 1 mM EDTA; 1% Triton X-100; 0.1% Deoxycholate, Na salt) with 1/100 PIC (Protease inhibitor cocktail Set IV, in DMSO; #539136, Merck) and 1 mM phenylmethanesulfonyl fluoride (PMSF, #P7626, Sigma-Aldrich). The cell suspension was divided into four 2-ml breaking tubes (BioSpec Products, #10832, Microtube 2 ml with cap) containing a 1-ml volume of glass beads (#11079105, BioSpec Products). Lysis was performed by ten cycles of 1-min beating, followed by 1-min chilling on ice (BioSpec Mini-BeadBeater-16, BioSpec Products). Cell lysates and glass beads were separated by punching a hole in the bottom of the tube with a red-hot 18G needle and then performing slow centrifugation (500 x *g*, 3 min, 4 °C; Eppendorf 5810R centrifuge, A-4-62 rotor). The collected lysates were combined into a 15-ml centrifuge tube and the pellet and supernatant were separated by centrifugation (12K rpm, 5 min, 4 °C; Eppendorf 5810R centrifuge, F-34-6-38 rotor). Pellets represent cell debris and the supernatant is the whole cell lysate, so the supernatant was collected and the final volume was made to 4 ml using FA buffer with 1/100 PIC and 1 mM PMSF in a new 15-ml centrifuge tube (#CFT011150, JET BIOFIL). About 40 μl of supernatant corresponding to 5 OD_600_ of cells was collected and stored at -20 °C as the input protein.

Immunoprecipitation was performed by incubating all of the lysate with 60 μl Anti-FLAG M2 Magnetic Beads (#M8823, affinity-isolated antibody, Merck) in 4 ml of FA buffer with 1/100 PIC and 1 mM PMSF. Additionally, MgCl_2_ and Benzonaze was added to keep the lysate stable. Binding was performed in a 15-ml tube with mixing using an end-over-end rotator (Intelli-Mixer, RM-2L, F1 mode, 30 rpm) at 4 °C for 16 h. The magnetic beads were anchored with a DynaMagTM-2 Magnet and the bound complexes were washed four times with 1 ml of each wash buffer for 5 min with mixing (Intelli-Mixer, RM-2L, F1 mode, 30 rpm) at room temperature. The bound complexes were eluted twice by heating at 65 °C in 100 μl 1X SDS sample buffer (0.065 M Tris-HCl, pH 6.8; 2% SDS; 10% Glycerol; 5% β-mercaptoethanol; 0.005% Bromophenol blue) for 20 min and, secondarily, in 100 μl 1X SDS sample buffer for 20 min. Two eluates were pooled as the Input-protein. Input-protein was diluted with 2X SDS sample buffer (0.125 M Tris-HCl, pH 6.8; 4% SDS; 20% Glycerol; 10% β-mercaptoethanol; 0.005% Bromophenol blue), heated at 65 °C for 20 min, and then stored at -80 °C. Before loading them in an SDS-PAGE gel, the samples were taken from -80 °C storage and heated at 65 °C for 20 min (Hsu et al., 2021).

The 8% SDS-PAGE was run with samples and a marker at constant 70V for 30 min and 150V for 1 hour. Protein was transferred from the gel to the PVDF membrane overnight. Monoclonal anti-FLAG M2 antibody (#F3165, Sigma Aldrich) and c-Myc Monoclonal Antibody (9E10) (#MA1-980, Invitrogen by Thermo Fisher Scientific) were used to identify Tfc3-3Flag and Tfc1-13Myc proteins, respectively. Anti-glucose-6-phosphate dehydrogenase (G-6-PDH) (#A9521, Sigma Aldrich) was used to identify endogenous G6PDH levels in whole cell lysates. Input-proteins do not show enrichment in endogenous controls, such as endogenous G6PDH, but the *Kl-TFC7* replacement line showed distinct endogenous G6PDH signal every time the experiment was performed, indicating physiological differences in the *Sc-TFC7* wild type and *Kl-TFC7* replacement lines.

## Data availability

The ChIP-sequencing data generated and analyzed in this study are available from NCBI under the accession number GSE287186.

## Notes

### Competing Interest Statement

The authors have declared no competing interest.

